# Population structure and association mapping for hundred seed weight in mungbean minicore

**DOI:** 10.1101/2021.05.04.442577

**Authors:** Muhammad Arslan Akhtar, Muhammad Aslam, Roland Schafleitner, Muhammad Ahsan, Ghulam Murtaza

**Affiliations:** Department of Plant Breeding and Genetics, University of Agriculture, Faisalabad, Pakistan; World Vegetable Center, Vegetable Diversity and Improvement, Tainan, Taiwan; Institute of Soil and Environmental Sciences, University of Agriculture, Faisalabad, Pakistan

**Keywords:** Mungbean, seed weight, genome-wide association mapping, multilocation analysis

## Abstract

Mungbean is an important legume rich in protein and carbohydrates. It is cultivated mostly in south to south east Asia and east Africa. One of the important yield contributing traits in mungbean is 100 seed weight. Variation for this trait is available in the mungbean germplasm, and to facilitate improvement of this trait through breeding, genome-wide association mapping for 100 seed weight was carried out in the World Vegetable Center mungbean minicore collection. A total of 24,870 single nucleotide polymorphic markers were tested for association with 100 see weight in plants grown in Pakistan, Bangladesh, Myanmar and Taiwan and candidate loci associated with seed weight on chromosomes 1, 4, 6, 8 and 9 were identified. None of the QTLs was stable across all environments, but loci on chromosomes 4 and 8 were significant in Pakistan and Bangladesh and other loci on chromosome 8 were significantly associated in plants grown in Myanmar and Taiwan.

## Introduction

Mungbean (*Vigna radiata* (L.) Wilczek), also known as green gram, is an important legume crop cultivated in south to south-east Asia and east Africa. Worldwide, it is currently cultivated on around 7.3 million hectares and produces about 5.3 million tons with an average yield of 0.73 tons per hectare. Major contributors towards mungbean production are India, China and Myanmar. The average yield of mungbean in Pakistan is comparable to the global average, but the overall production is less because of the small area under cultivation. The short vegetative period, the ability to withstand drought stress and fix atmospheric nitrogen along with high nutrient content makes mungbean a popular legume crop. 100 grams of boiled mungbean grains contain 7.02 g protein, 19.15 g carbohydrates, 7.60 g dietary fiber and 0.40 g fats, whereas 22% vitamin B1, 80% vitamin B9, 24% Mg, 16% Fe and 11% Zn on the basis of daily nutrient requirement [1]. Apart from grain, other plant parts are used as fodder and manure. The short cropping duration makes mungbean fit for many cropping patterns and combinations.

Large seeded mungbean achieves premium prices on markets, therefore large 100 seed weight is often preferred. Information on genetic loci contributing to this trait across environments would enhance breeding for larger grain. Molecular markers linked to various traits can help in the screening and selection of parents for the development of high yielding and disease resistant mungbean genotypes [2]. Loci associated with seed size in mungbean have been mapped previously in bi-parental populations [3,4]. Genome-wide association mapping or genome wide association studies (GWAS) is a complementary genetic approach used to associate variations in phenotype in germplasm panels, testing more alleles than in bi-parental populations. In the present study, association between 100 seed weight and a genome-wide marker set was tested in a mungbean minicore collection at four locations in Pakistan (2016), Bangladesh (2017), Myanmar (2016) and Taiwan (2018).

## Materials and methods

A total of 293 mungbean genotypes from the minicore set were cultivated in Pakistan (Islamabad, 33°40’42.6”N, 73°08’20.4” E, from March to May, 2016), Bangladesh (Ishurdi, 24°07’43”N, 89°03’57” E, from March to May 2017)Myanmar (Yezin, 19°50’02” N, 96°16’45” E, 133 m above sea level, from November 2016 to January 2017), and in Taiwan (World Vegetable Center, 23°01’30” N, 120°17’35” E, from September to November 2018). Data for 100 seed weight were determined on 3 replications of 5 plants per accession.

Genotyping data of the mungbean minicore accessions (Table 1) were obtained from Dart PRL as described by our fellows [5]. Imputation for the missing data was done using LD KNNi imputation protocol [6]. SNPs with more than 10% missing data after imputation and less than 5% minor allele frequency were filtered out.

**Table 1.**
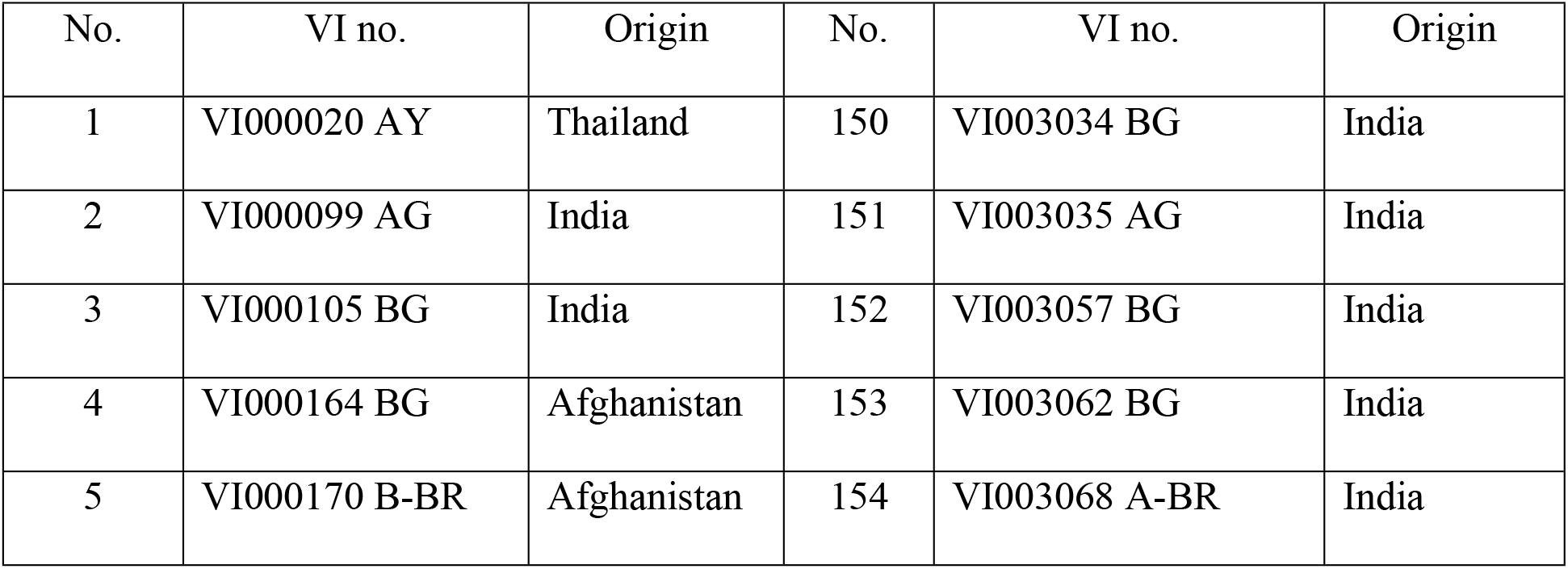
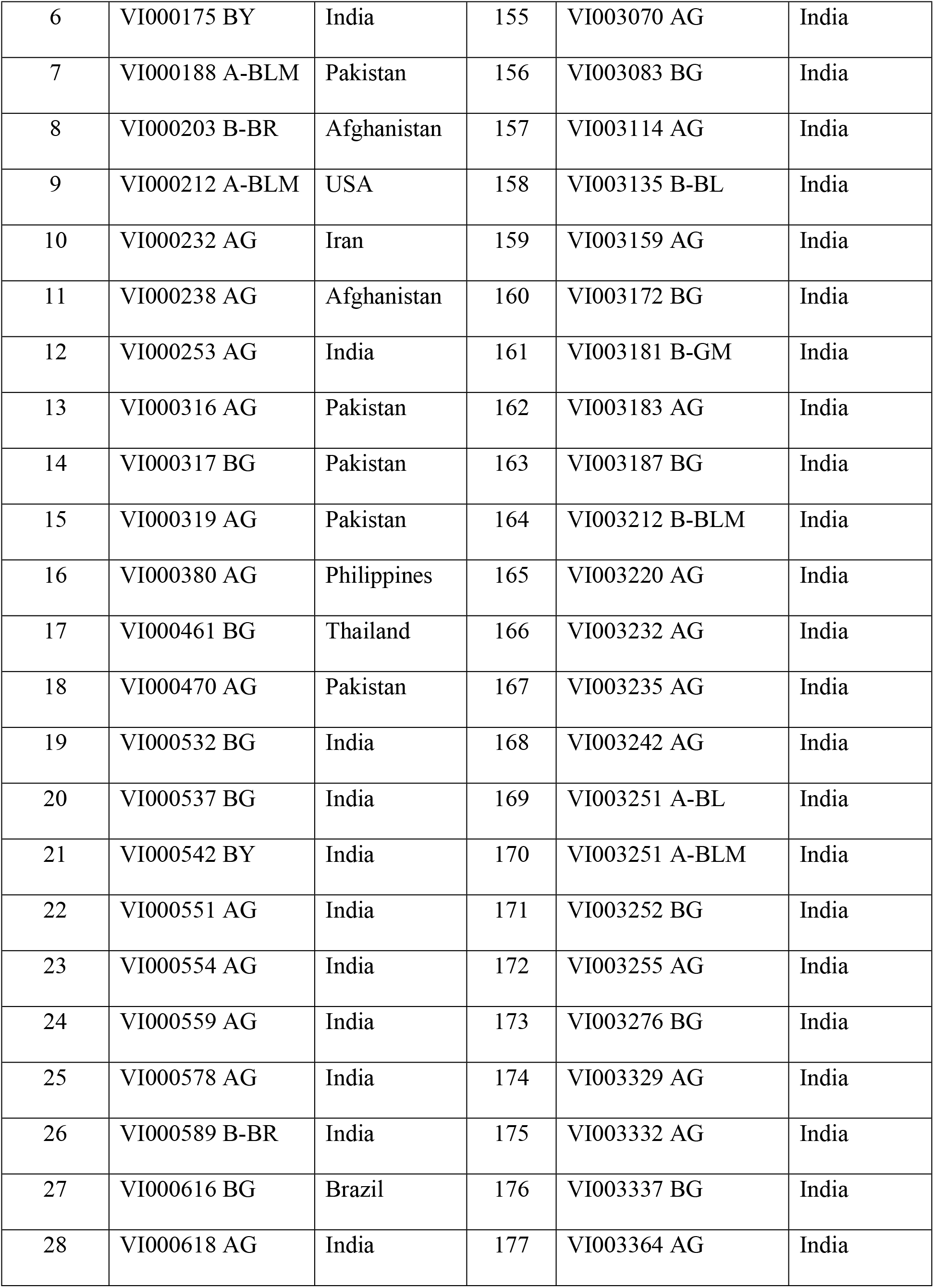
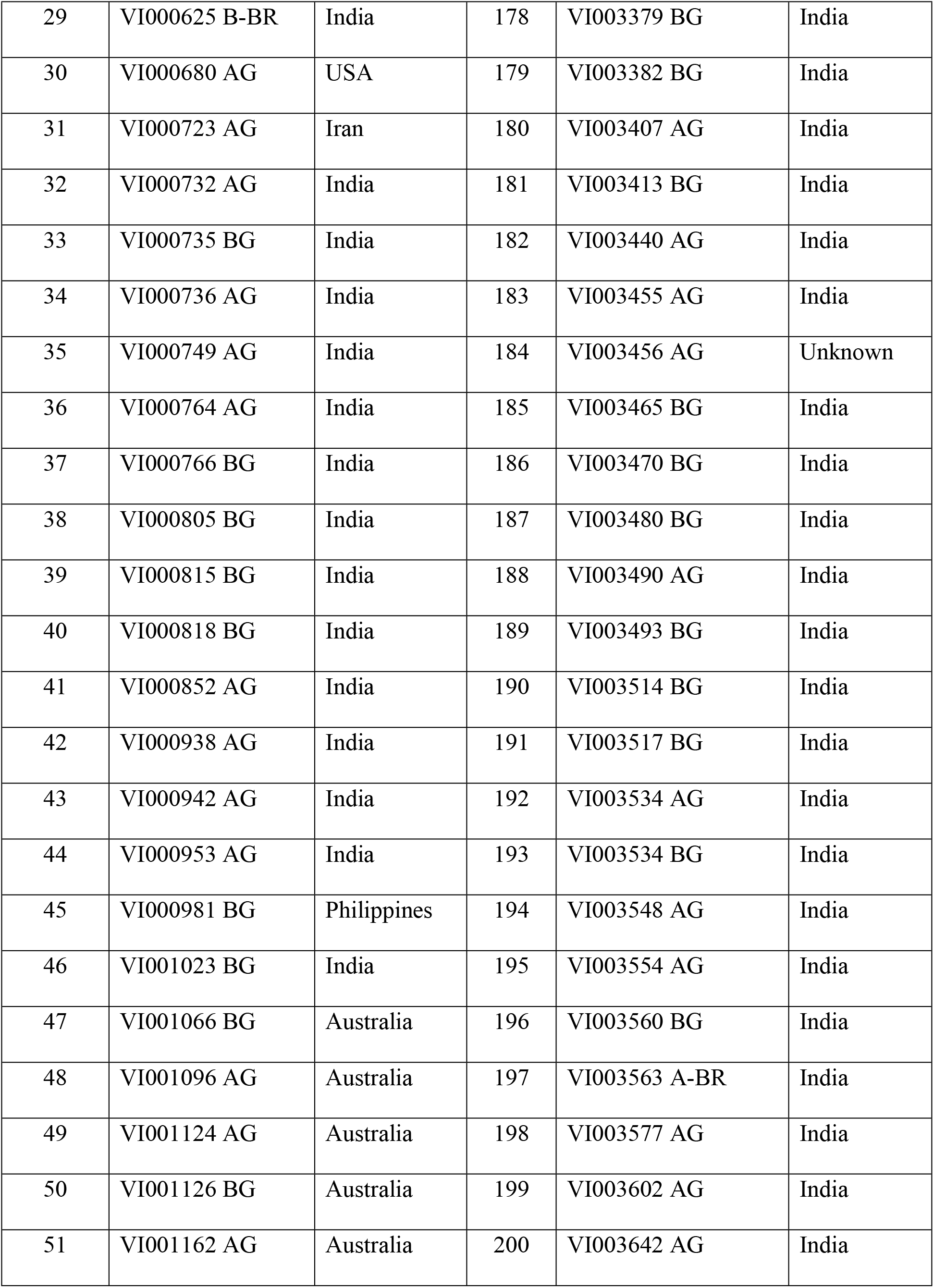
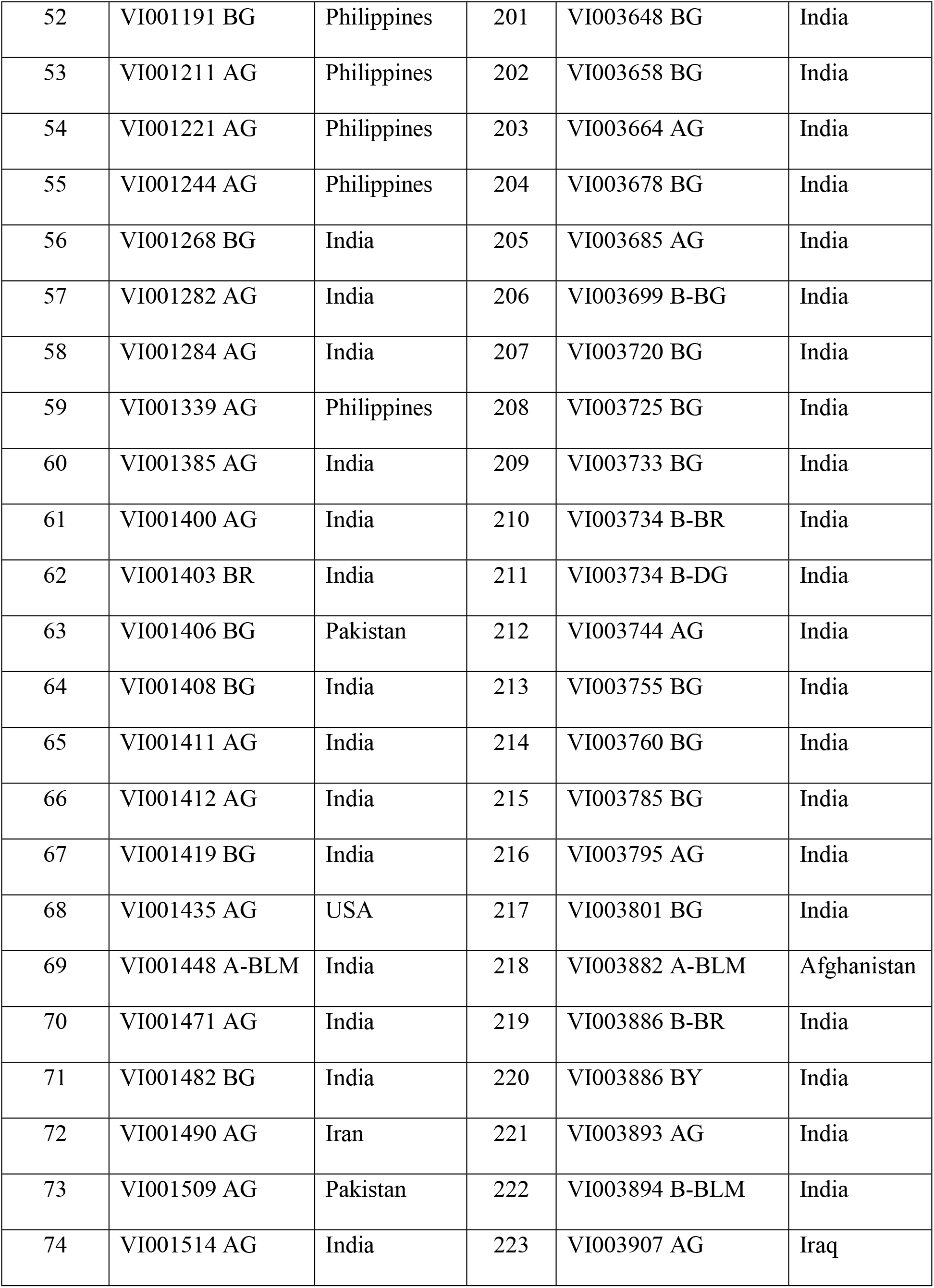
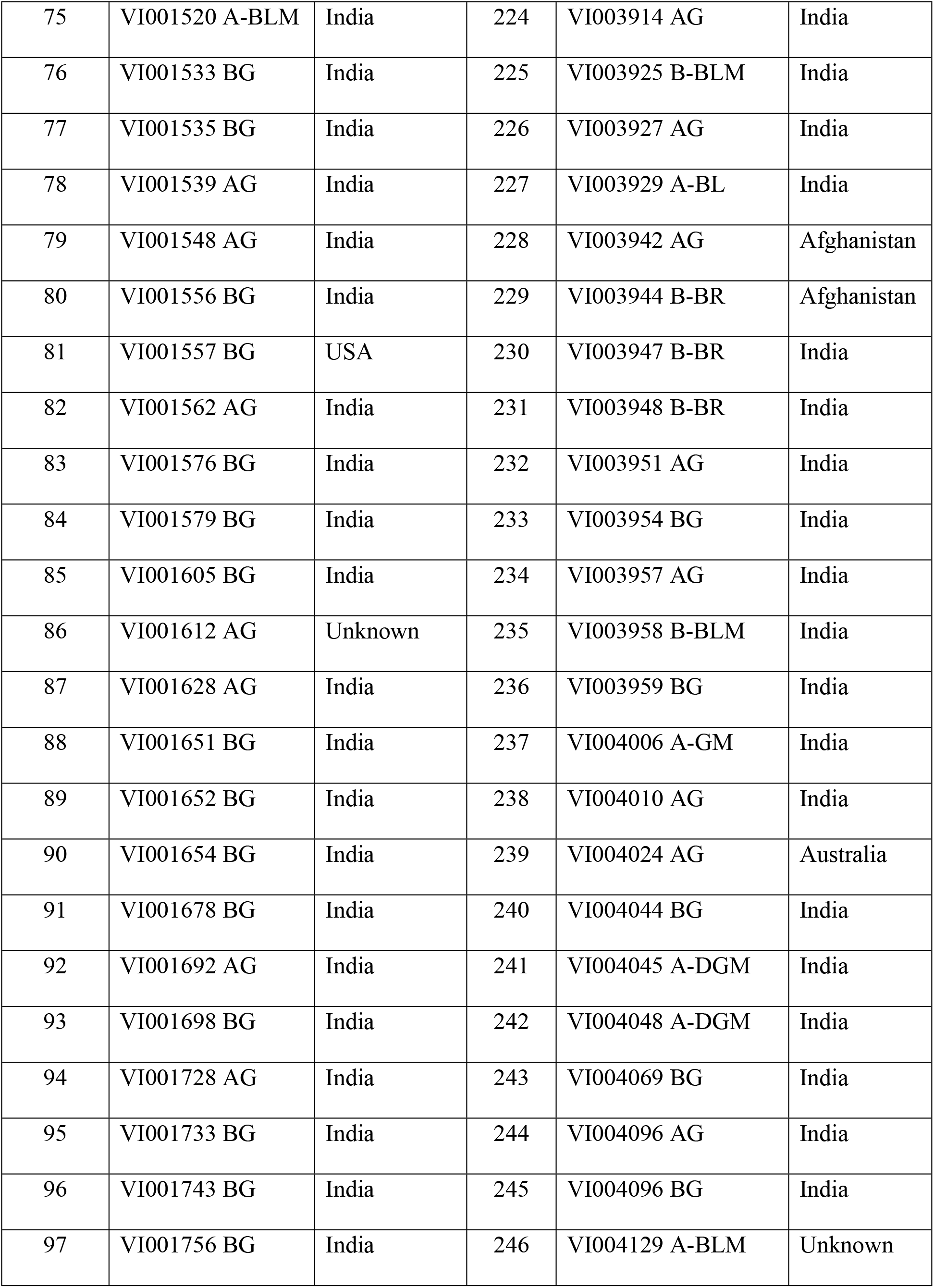
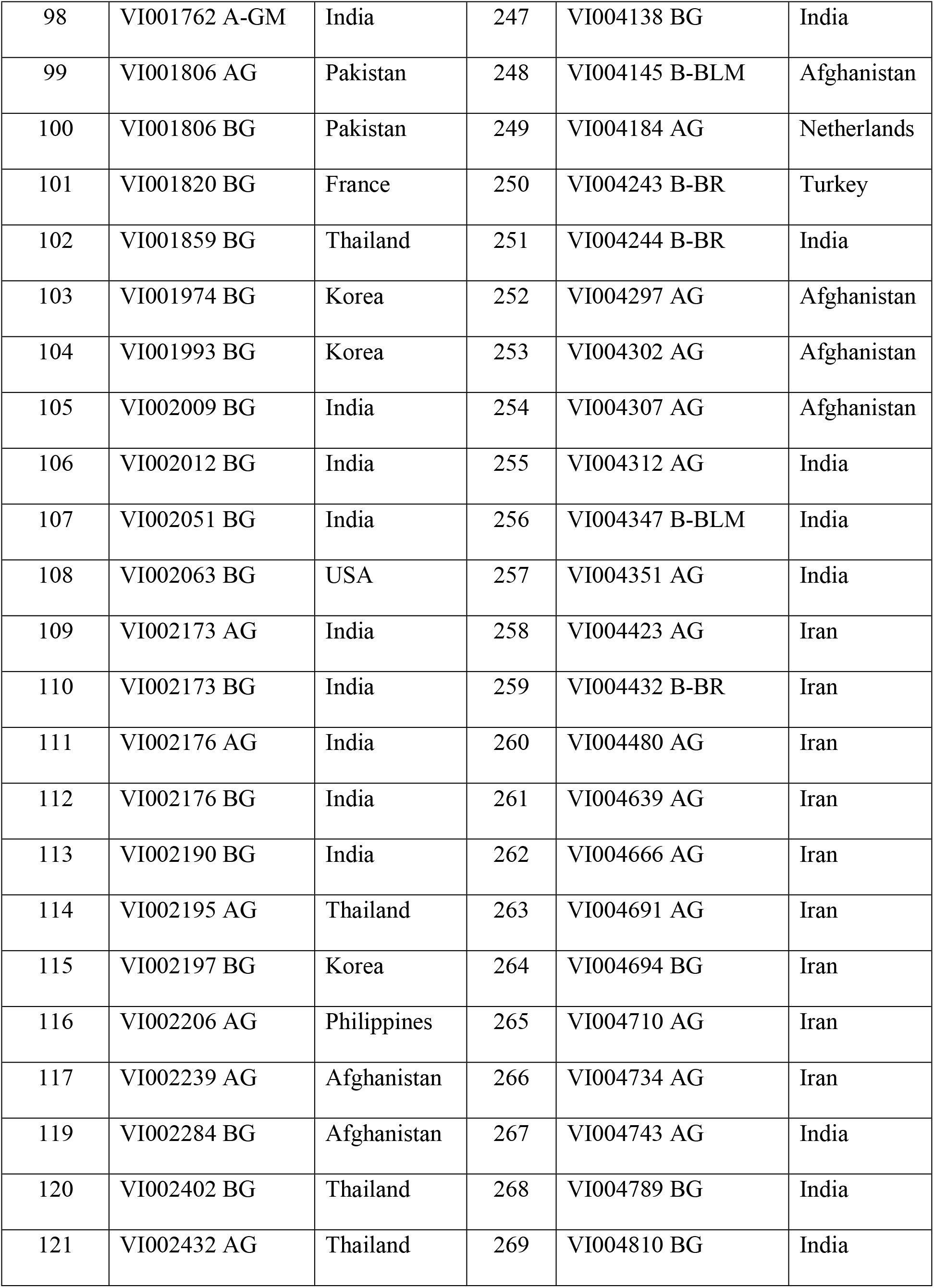
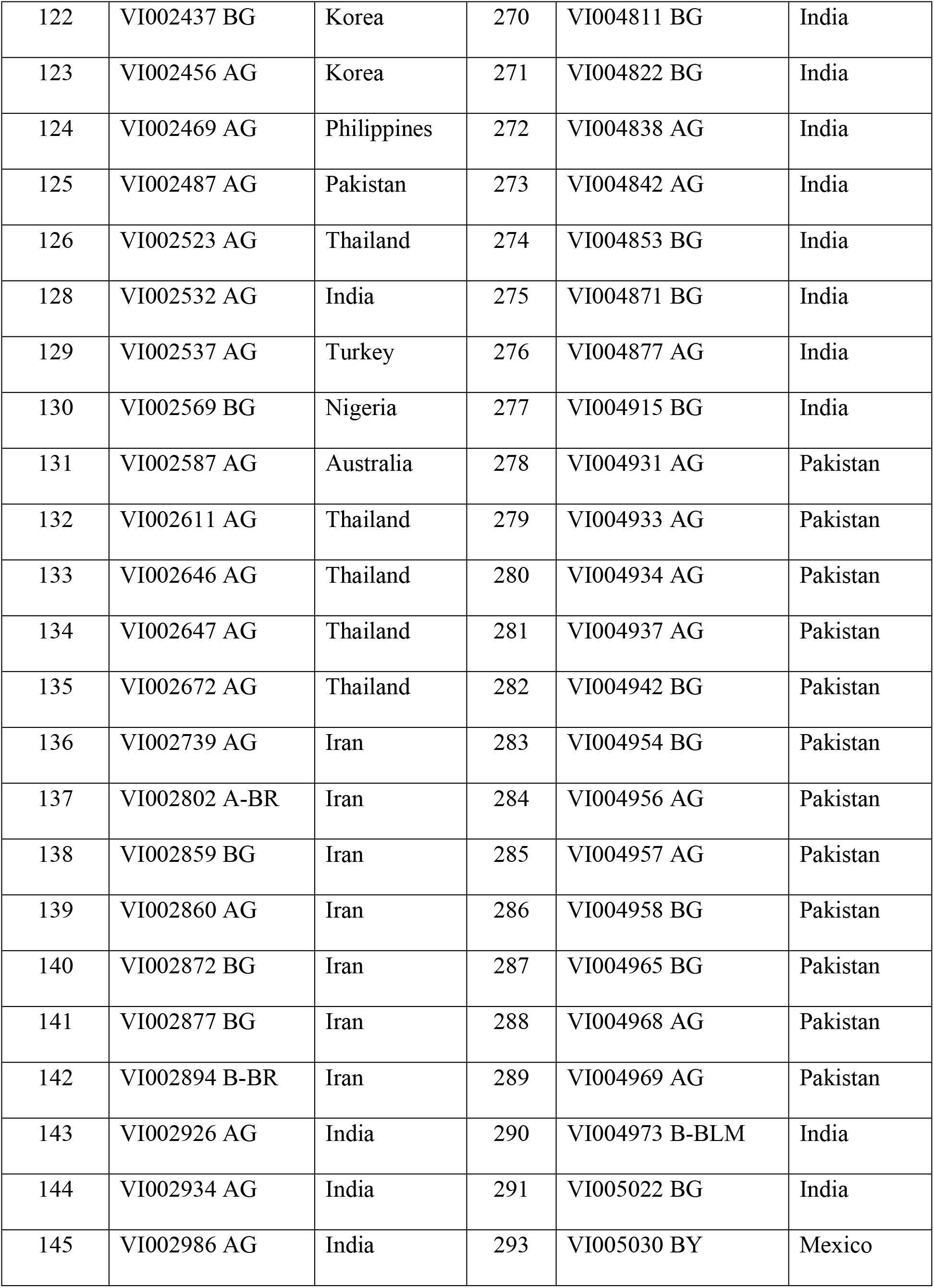
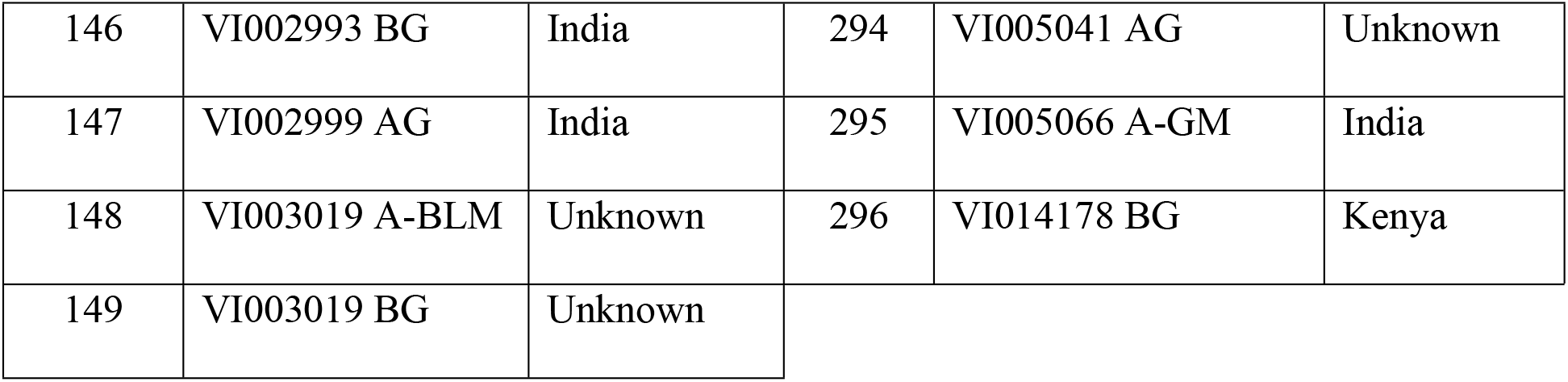
Mungbean minicore genotypes used under study.

Archaeopteryx tree was developed using complete SNP array of the minicore set following clustering method of neighbor joining [7] within TASSEL (Trait Analysis by aSSociation, Evolution and Linkage, version 5.0). Population structure analysis to be used as co-factor in GWAS was performed in STRUCTURE (version 2.3.4) to assess the population subgroups (K) using a Markov Chain Monte Carlo (MCMC) approach with a burn in period of 50,000 and 500,000 MCMC repeats for K values K = 2 to K = 15 [8,9]. The output was analyzed according to [10] using STRUCTURE HARVESTER [11], to identify the most likely population structure (K-value) for the germplasm panel under investigation. GWAS was performed in TASSEL. Candidate SNPs were first manually examined by comparing the phenotypic values with the marker alleles in Microsoft Excel.

## Results and discussion

The 100 SW data obtained from 4 environments were compared. Overall, seed size among the locations was different, but correlated (Table 2, Fig 1). Seed size in average was largest in Pakistan and smallest in Bangladesh. Stability of 100 SW across environments was strongly different among the genotypes.

**Table 2.** Pearson correlation coefficients and T-test results of the comparison of 100 SW across 4 locations in Bangladesh, Pakistan, Taiwan and Myanmar.

|  |  |  |  |  |
| --- | --- | --- | --- | --- |
| Pearson |  | Pakistan | Taiwan | Myanmar |
|  | Bangladesh | 0.48<br>(p<0.001) | 0.6<br>(p<0.001) | 0.56 (p<0.001) |
|  | Pakistan |  | 0.62<br>(p<0.001) | 0.65 (p<0.001) |
|  | Taiwan |  |  | 0.81 (p<0.001) |
| T-test |  | Pakistan | Taiwan | Myanmar |
| (p-value) | Bangladesh | 7.59 E-83 | 0.01 | 0.00003 |
|  | Pakistan |  | 3.9 E-81 | 2.52 E-55 |

**Figure 1.**
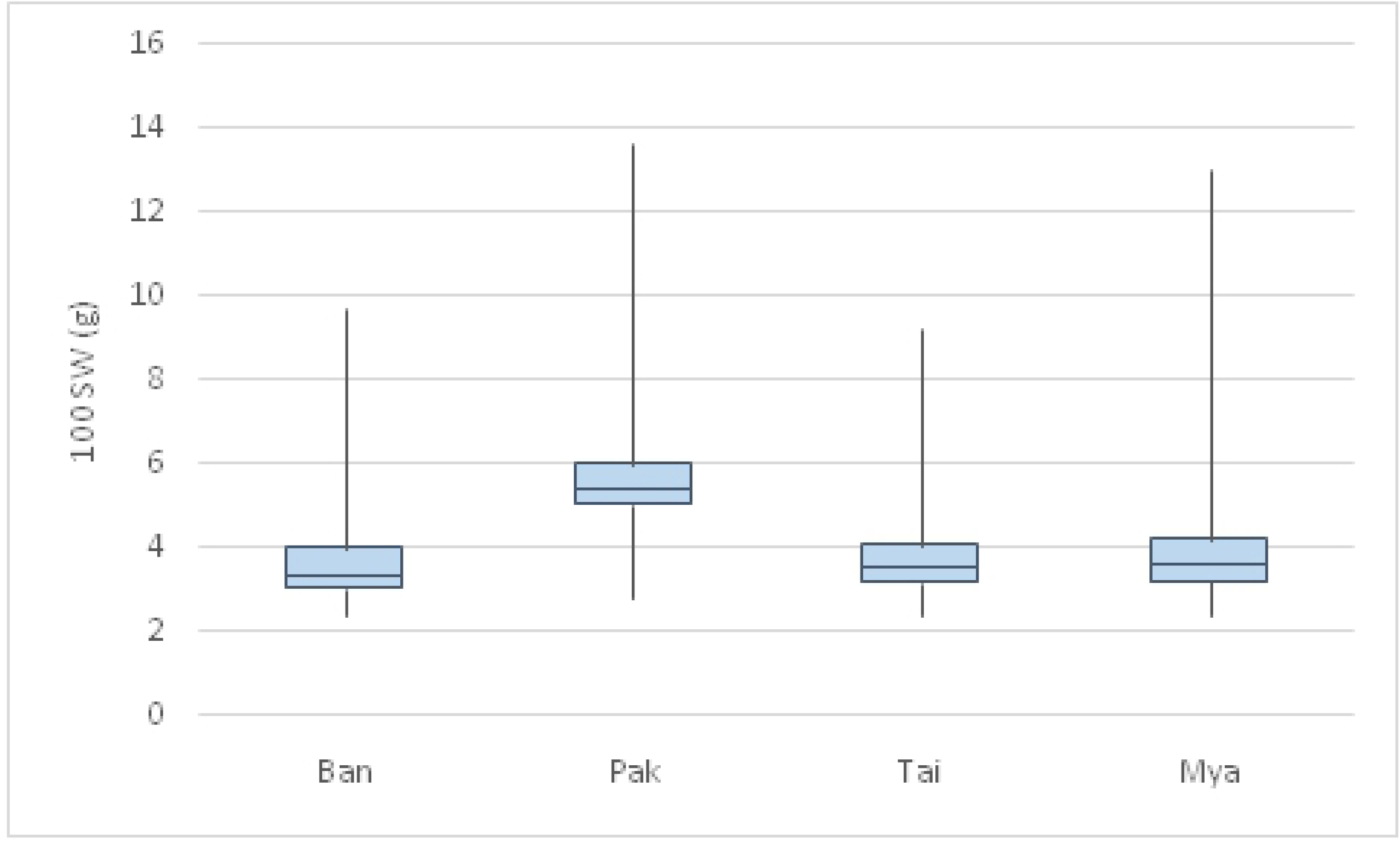

The minicore set was genotyped with 6,506 SNP markers distributed over all 11 chromosomes. Population structure analysis indicated that the minicore collection contained 3 sub-populations. L(K) graph (Figs 2a and 2b) shows sharp variation from cluster 2 to cluster 3, and delta K (ΔK) showing peak at cluster 3 indicates that the genotypes under study belong to 3 main clusters contrary to the findings of [5] who found out number of possible sub-populations to be 2 or 4. Seed weight was not evenly distributed over the subpopulations. In average, 100SW was larger in subpopulation 2 (average 5.9 g) than in subpopulation 1 and 2 (average 3.8 g). The size differences between subpopulation 2 and the other two subpopulations was significant (p< 0.001).

**Figure 2a.**
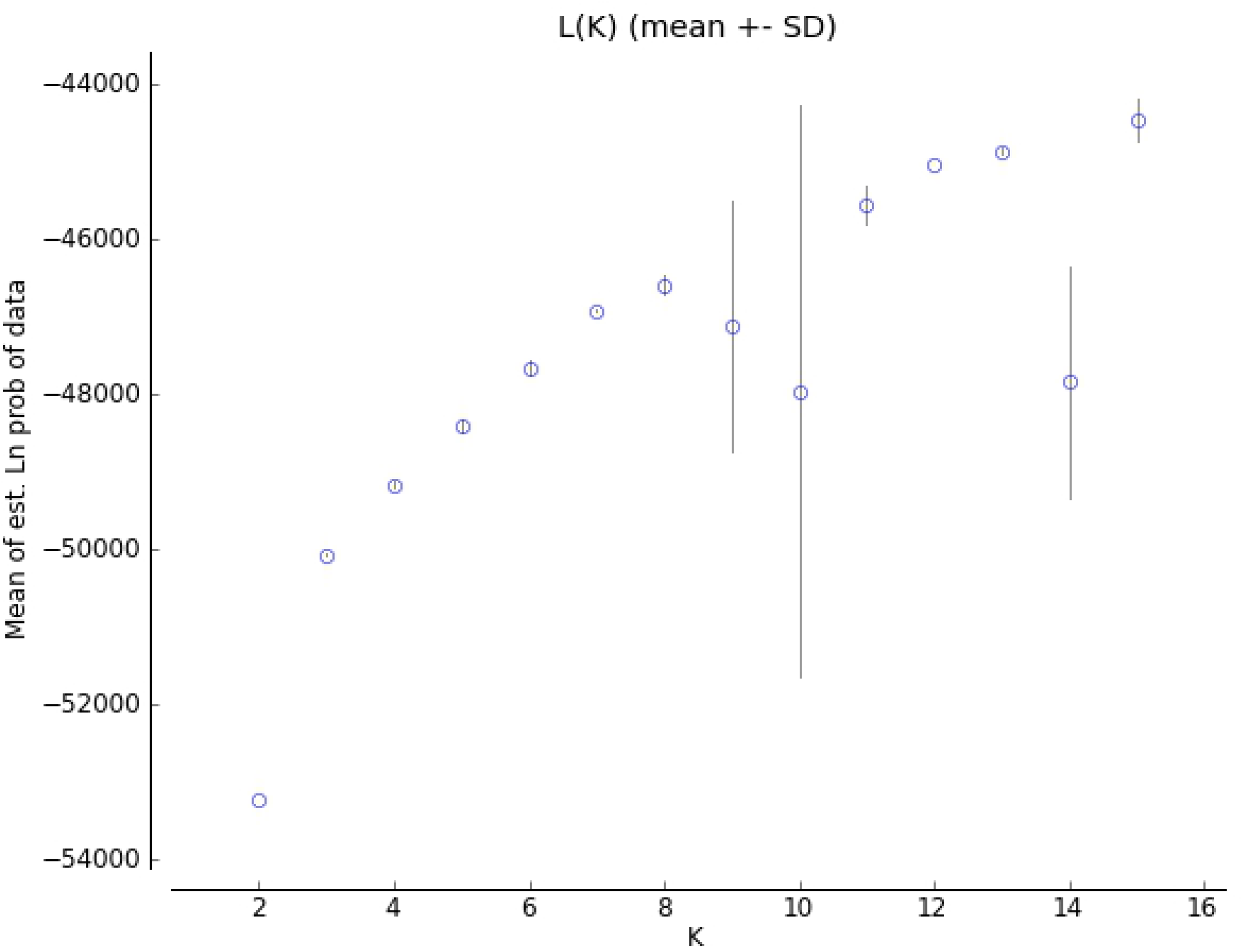

**Figure 2b.**
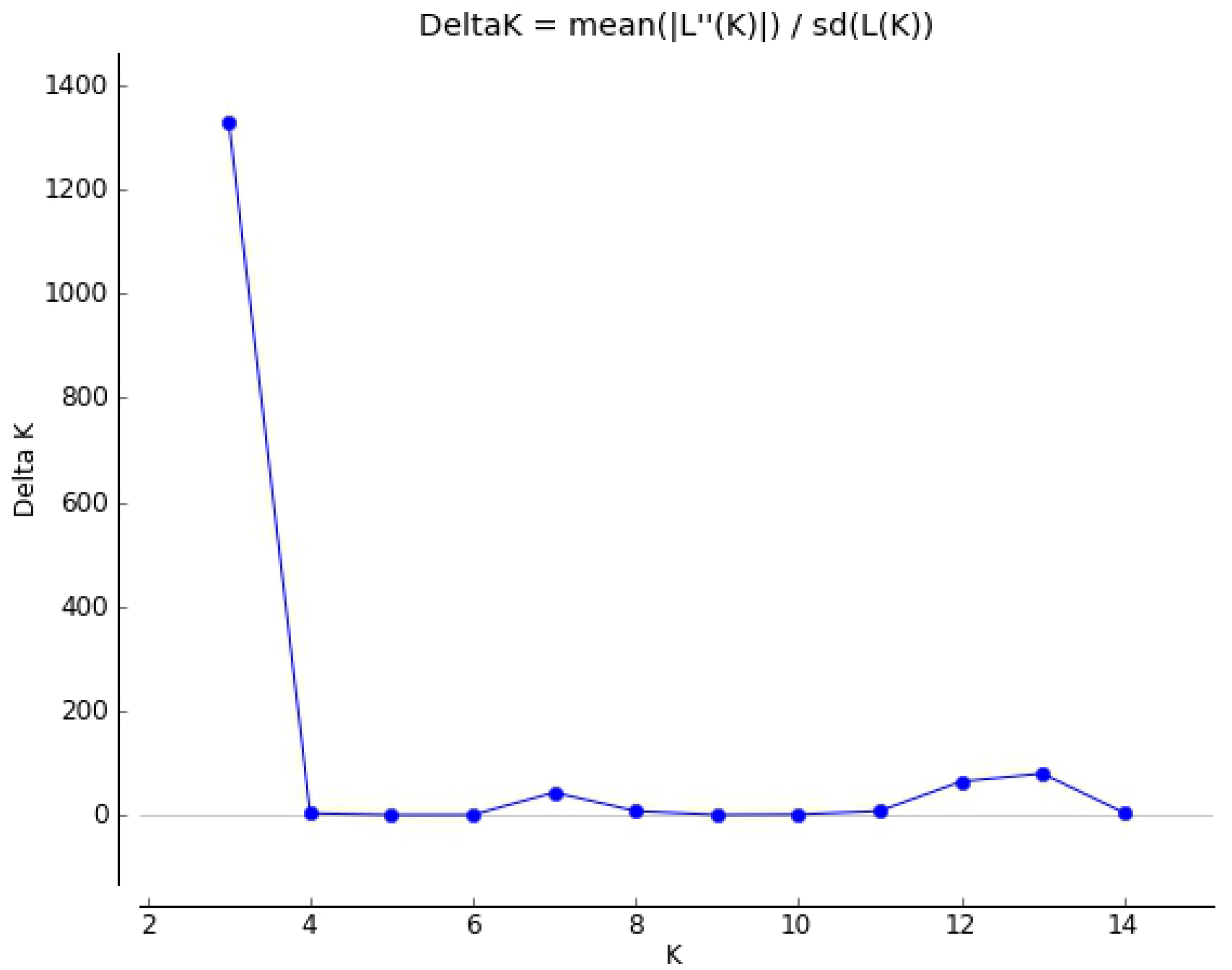

Population structure graph developed by STRUCTURE (Fig 3) showed the distribution of the genotypes under study in their respective sub-populations. Most of the accessions showed at least some degree of admixture.

**Figure 3.**
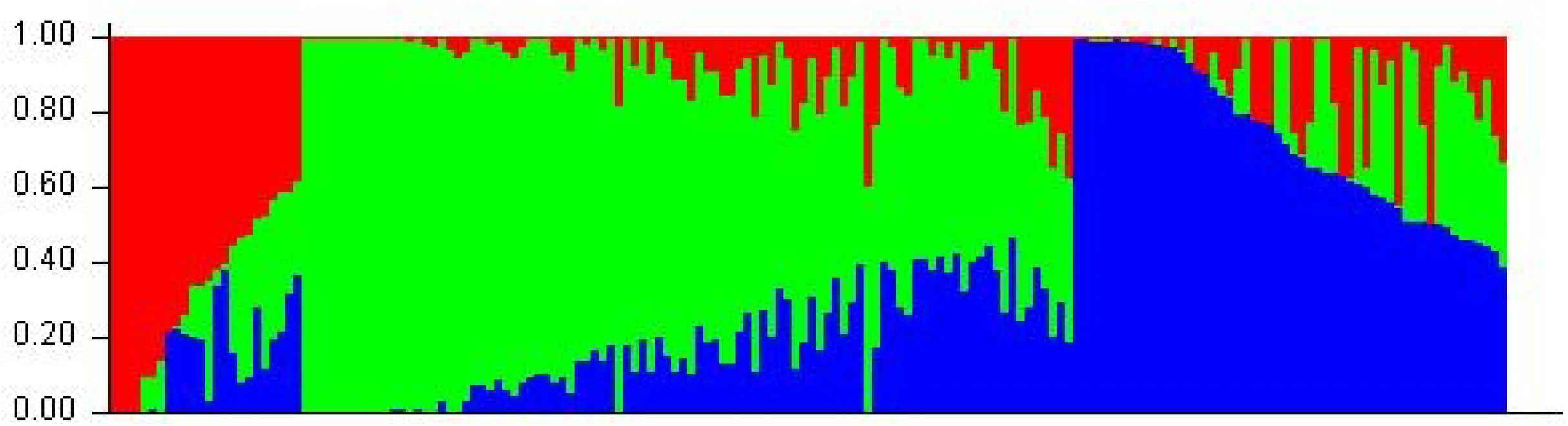

The population structure was in line with a phylogenetic tree That contained 3 major clusters, divided into sub-clusters (Fig 4). Quantile-quantile (QQ) graphs plot significance of associated SNP markers to the studied trait. The inclined ascending line shows the expected probability of the association of SNPs assuming there is no association of the trait of interest with SNPs studied, while the dots around the line show actual calculated probabilities. Deviation from the expected probability indicates association between SNPs and the trait of interest. The greater the difference between the expected line and calculated probability, the stronger will be the association [12]. QQ plots from all four locations showed different levels of association. The largest number of associated SNPs was obtained for the trial in case Myanmar, followed by Bangladesh, Pakistan and Taiwan.

**Figure 4.**
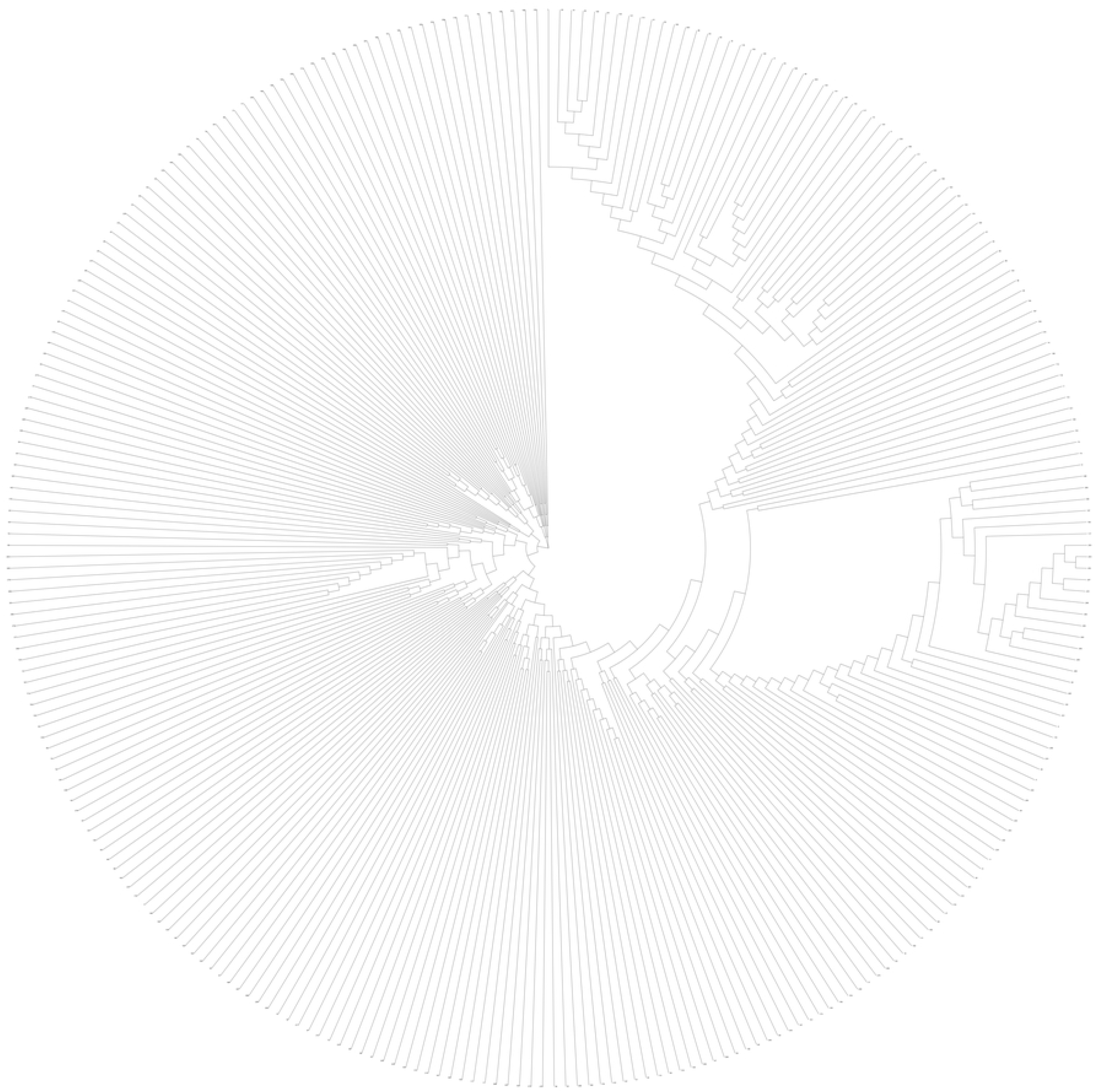

Manhattan plots were developed to depict the association of SNP positions with 100 seed weight (Fig 6). Random dispersion of the SNPs indicates that there is no significant association, while dots above a certain threshold, here assumed as 4.0 (-log10 of p-value) indicate significant association. Vertical alignment of associated SNPs is formed if linked SNPs at a locus are significantly associated with the trait. This facilitates the identification of nearby genes as candidates for being involved with the trait variation. The analysis of the marker-trait association from Pakistan showed vertically aligned markers in chromosomes 2, 7 and 8 (Fig 6a) pinpointing loci associated with 100 SW variation. In case of Bangladesh and Taiwan, aligned markers were present at chromosome 6, 7 and 8 (Figs 6b and 6d), and for Myanmar candidate markers were found on chromosomes 7 and 8 (Fig 6c).

**Figure 5a.**
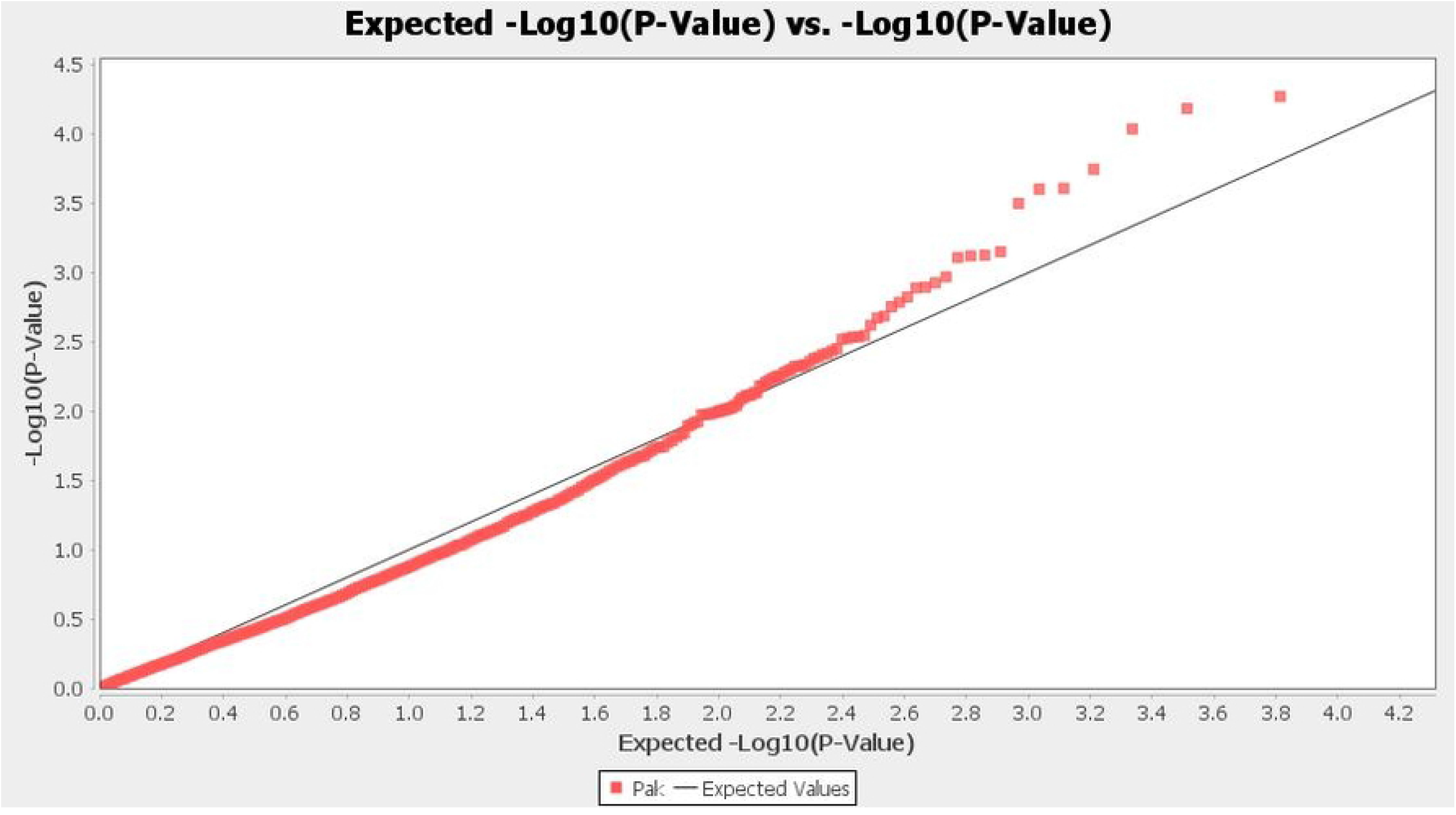

**Figure 5b.**
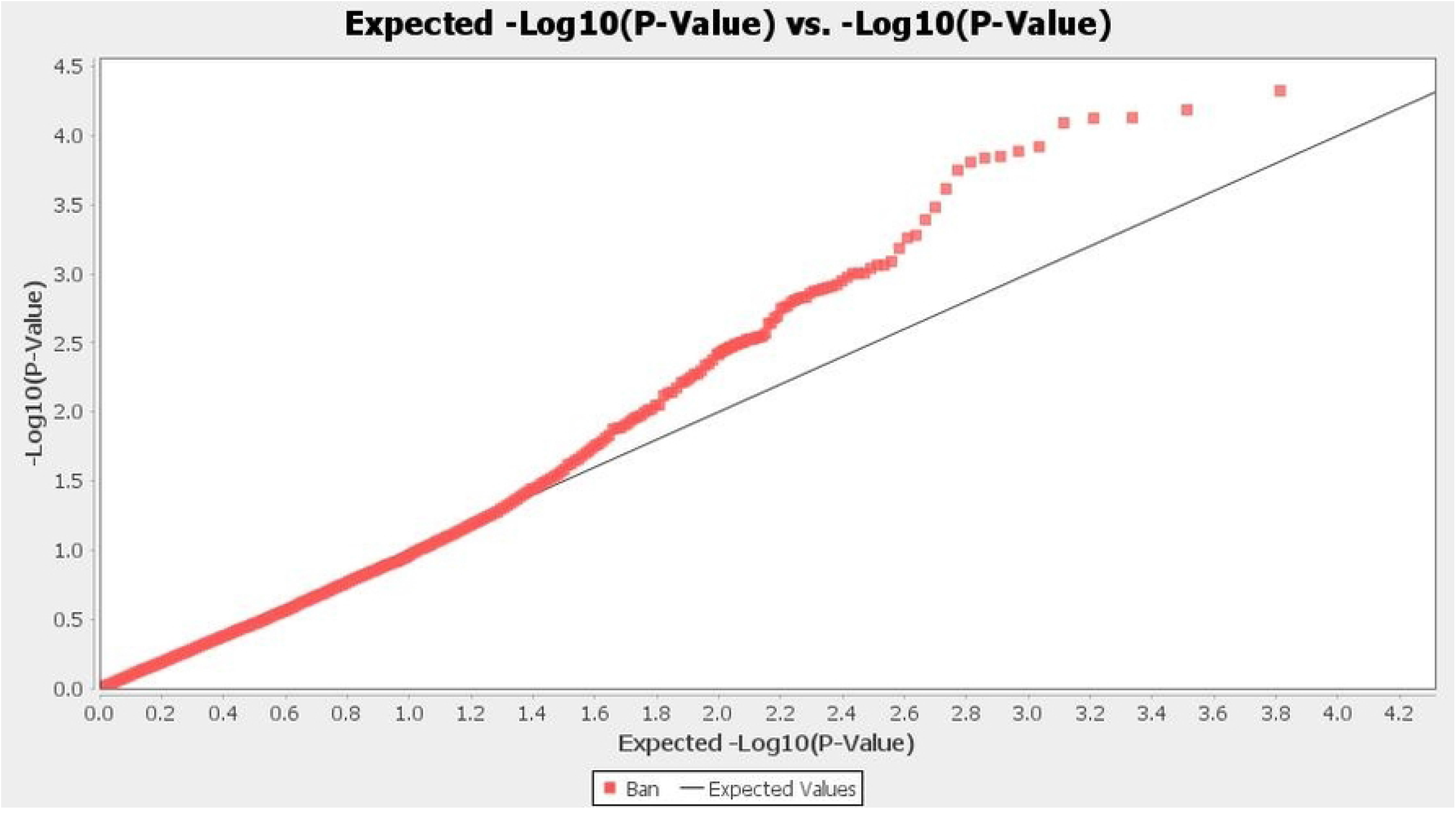

**Figure 5c.**
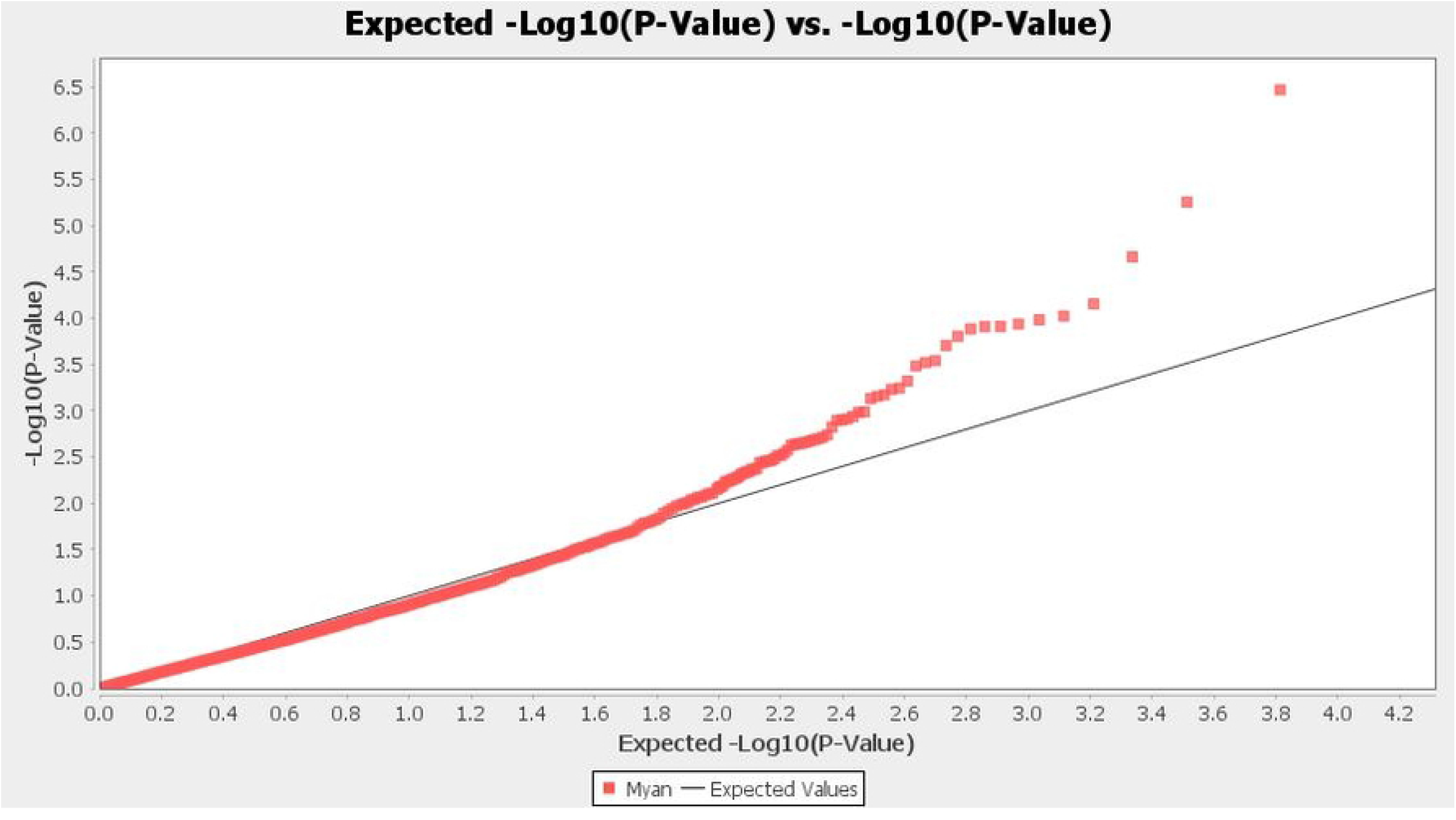

**Figure 5d.**
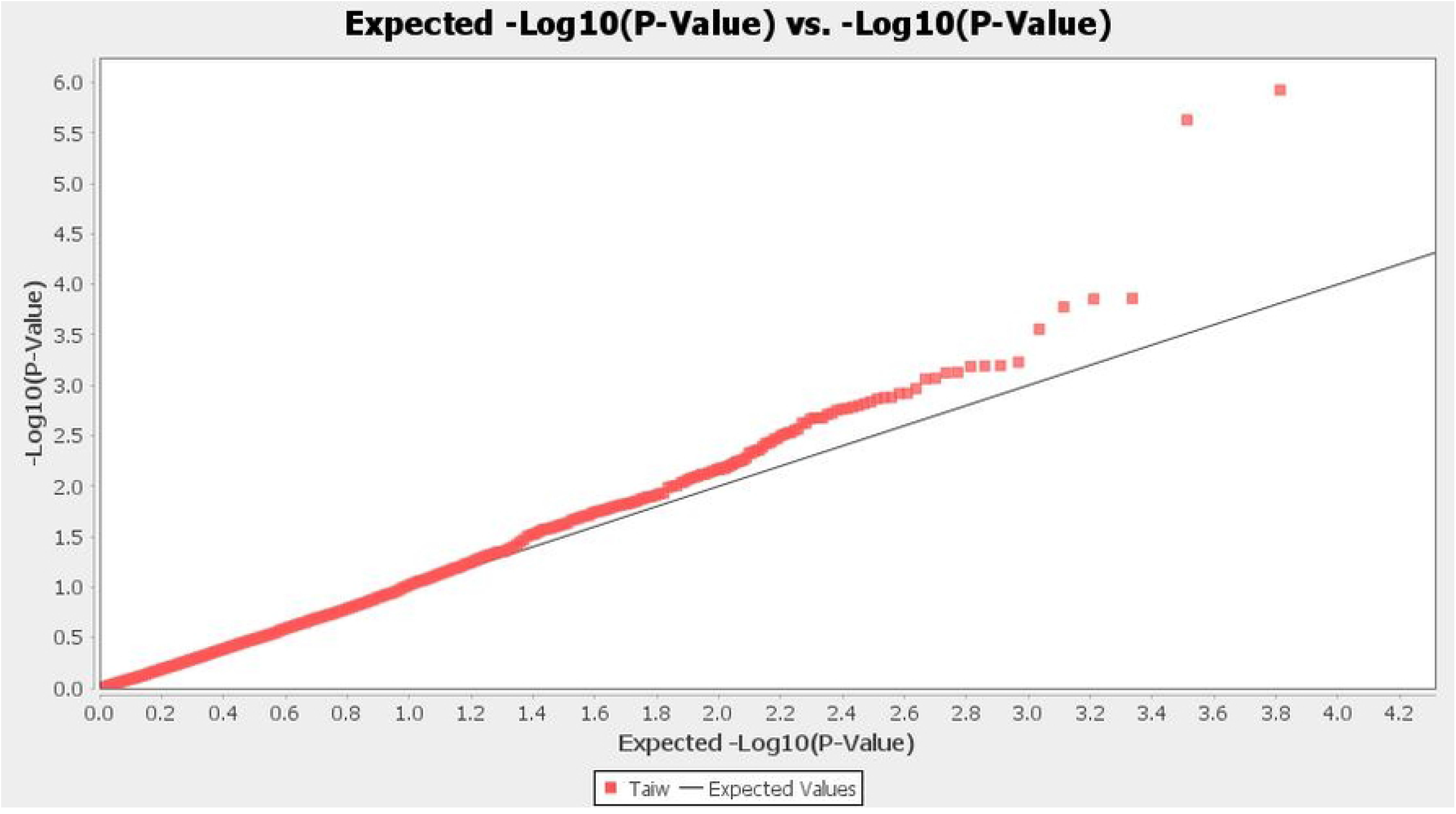

**Figure 6a.**
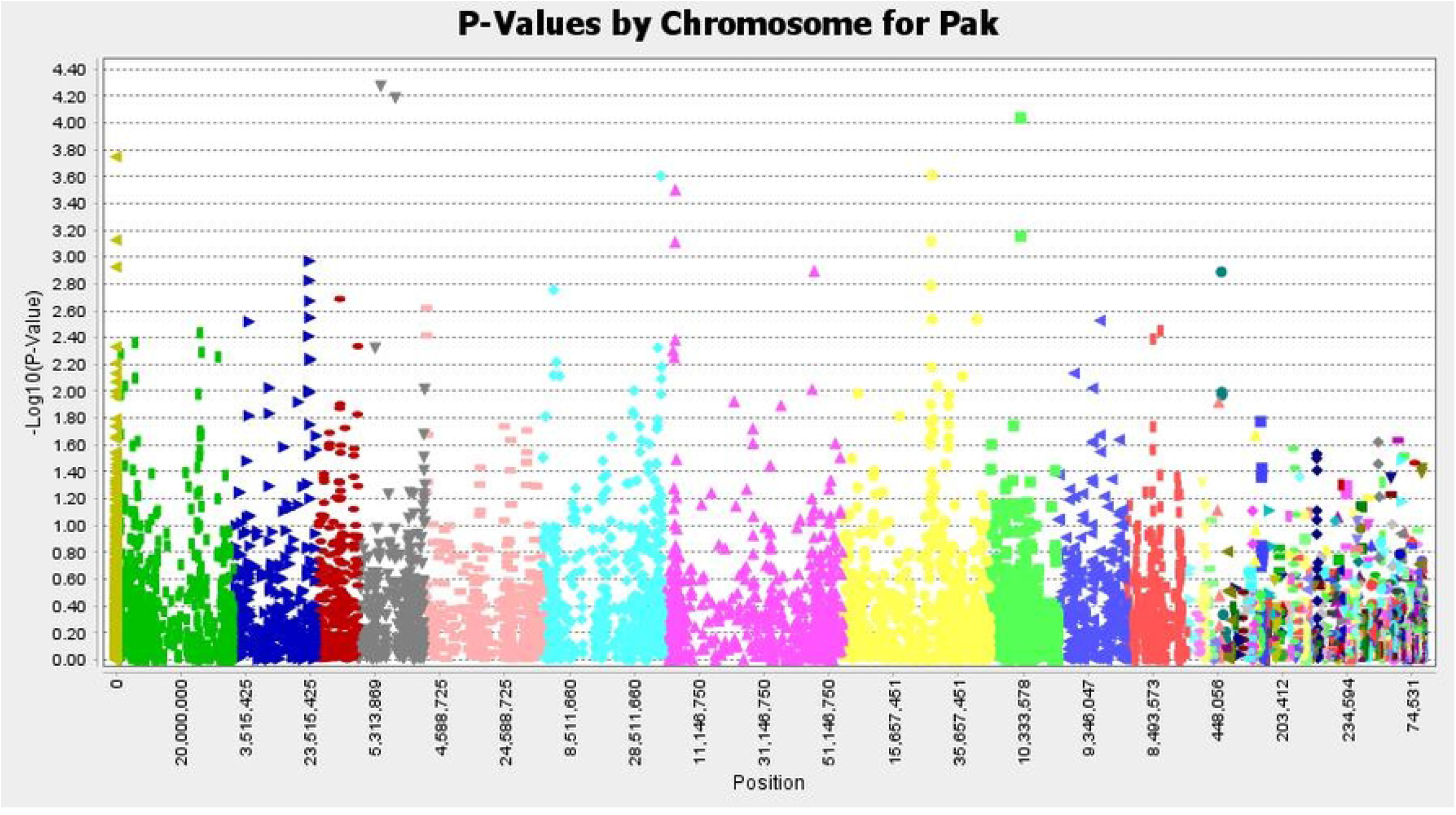

**Figure 6b.**
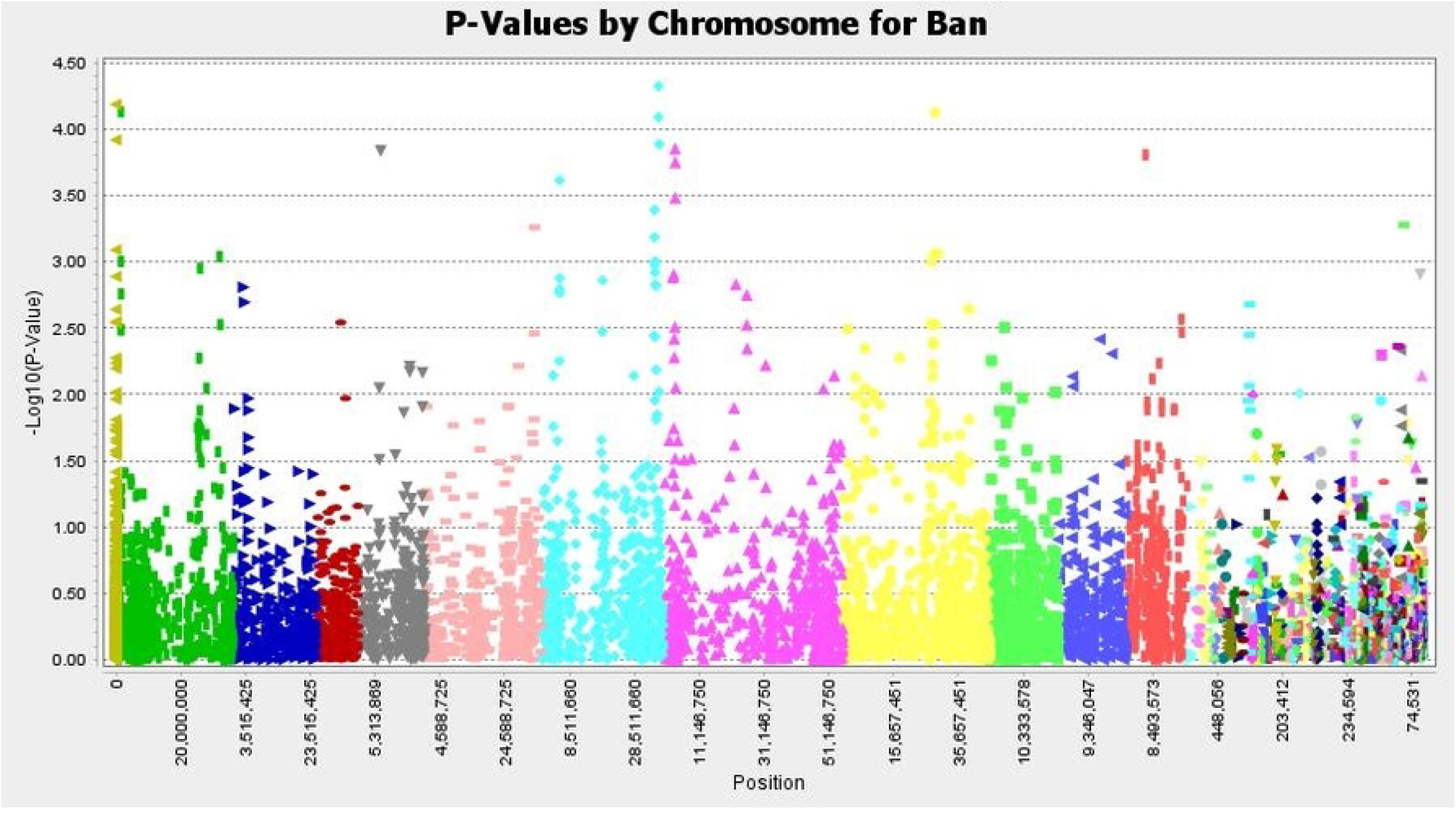

**Figure 6c.**
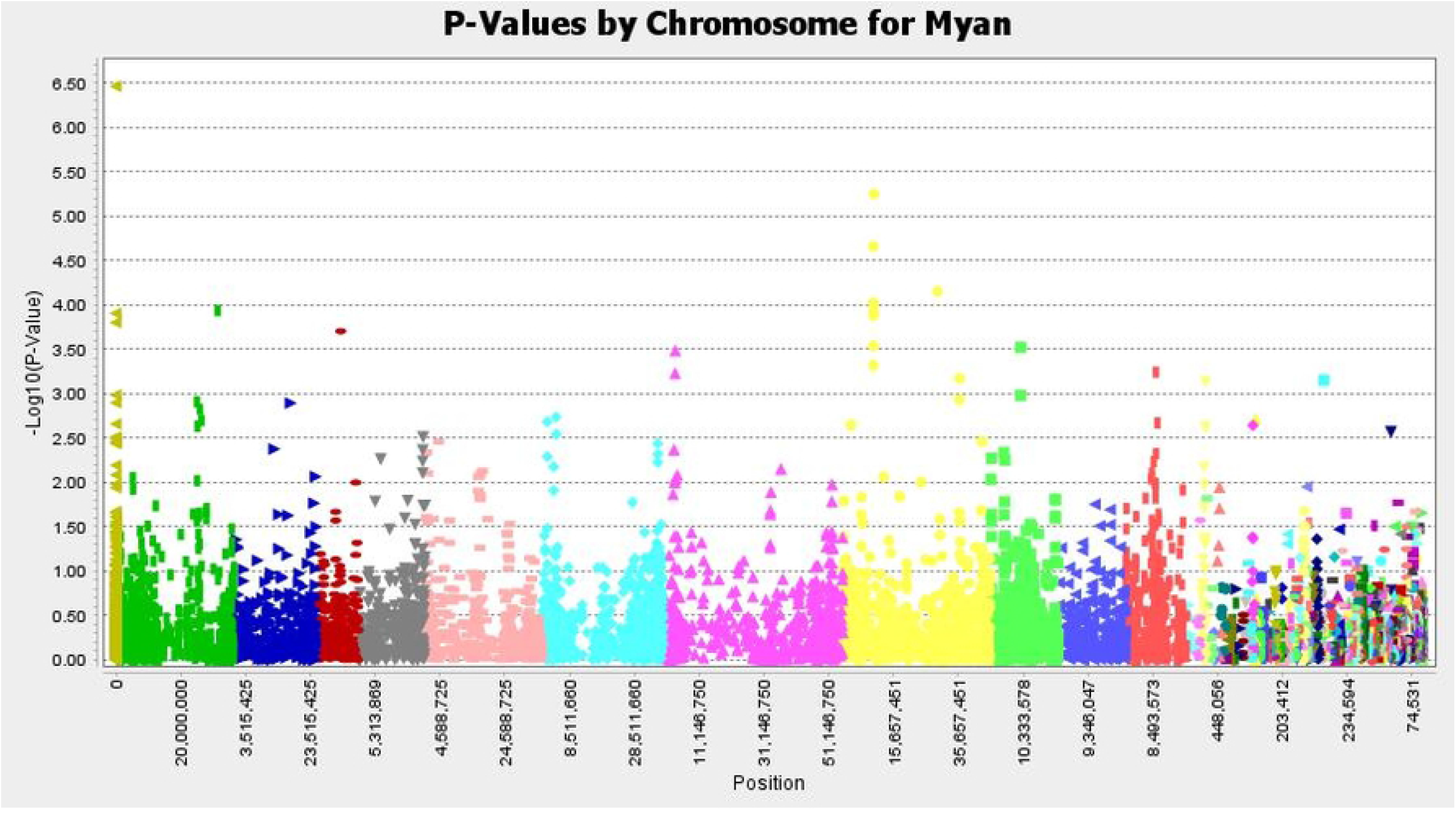

**Figure 6d.**
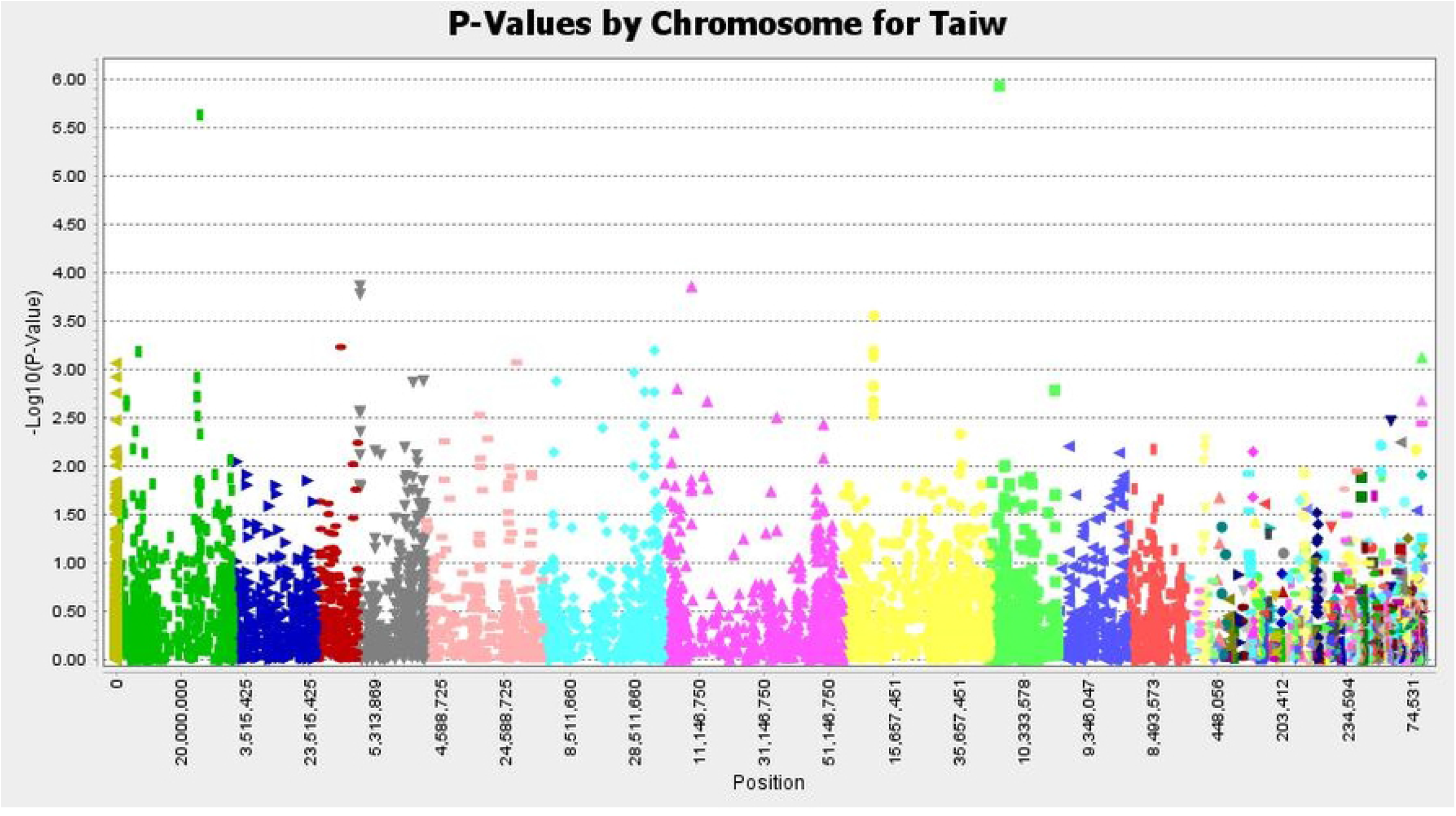

The candidate SNPs were aligned along with genotypes and 100 seed weight arranged in ascending order to depict the association between SNP genotype and increased 100 seed weight. This helped in the screening of the false positive markers and selection of markers that were showing some degree of relatedness. It can be seen from table 2 that most of the associated sites are located on chromosome 8. Additionally, some sites are also commonly found associated with 100 seed weight across multiple locations. It means that the gene(s) for 100 seed weight is possibly locate around said positions of the markers on chromosome 8 (Table 3).

**Table 3.** SNPs showing significant association with 100SW across locations.

| Chromosome | Pakistan<br>SNP position | Bangladesh<br>SNP position | Myanmar<br>SNP position | Taiwan<br>SNP position |
| --- | --- | --- | --- | --- |
| 1 |  | 32,026,195 | 31,425,058 | 25,049,038 |
| 4 | 7,032,154 | 7,032,154 |  |  |
|  | 11,559,392 |  |  |  |
| 6 |  | 5,433,578<br>34,701,957<br>34,712,555<br>34,744,525<br>34,913,611<br>35,030,031 |  |  |
|  | 36,662,459 |  |  |  |
| 8 |  |  | 9,321,413<br>9,379,508<br>9,427,417<br>9,427,833 |  |
|  |  |  | 9,427,846 | 9,427,846 |
|  |  |  | 9,432,667 | 9,432,667 |
|  |  |  | 9,588,677 | 9,588,677 |
|  | 27,458,129 |  |  |  |
|  | 27,246,365 | 27,246,365 |  |  |
|  |  | 28,480,139 |  |  |
|  |  | 28,480,205 |  |  |
|  |  |  | 29,255,058 |  |
|  |  |  | 35,992,362 |  |
|  |  |  | 35,978,574 |  |
| 9 | 9,495,582 |  |  |  |
|  | 9,488,553 |  | 9,488,553 |  |

Genetic analysis of domestication related traits in a mungbean population derived from the wild accession JP211874 collected in Myanmar and the mungbean cultivar Sukhothai pointed towards loci on linkage group 8 (corresponding to chromosome 8 of the reference sequence [13] between markers CEDG059 (position ∼ 15,462,360 bp) and VM37 (position ∼29,308.079) [3] overlapping with markers found associated with seed size in Pakistan and Bangladesh. The seed size QTLs mapped by [4] could not be attributed to specific chromosomes as for most associated markers nor sequence was disclosed that could be attributed to a physical position.

Association of markers to 100 SW was mostly specific for a specific trial. Only a few loci on chromosome 8 were significant in trials in Myanmar and Taiwan, or between Pakistan and Bangladesh. The trial in Myanmar was performed in the monsoon season, while the trial in Taiwan was planted in the pre-monsoon season.

## Conclusion

Mungbean minicore set used in present study splits into 3 sub-populations. Most of the markers putatively associated to 100 seed weight in mungbean are located on chromosome 8. The association between SW and the marker at position 8:27,246,365 bp was common for the trials in Pakistan and Bangladesh data, whereas markers 8:9,588,667, 8:9,427,846 and 9,432,667 were common in trials in Myanmar and Taiwan. These markers are likely associated with 100 seed weight in mungbean and further investigations and validations are needed to confirm these results.

## Acknowledgement

The above research was conducted at the World Vegetable Center, Taiwan, and was funded by Higher Education Commission (HEC) of Pakistan under International Research Support Initiative Program (IRSIP) and by the long-term strategic donors to the World Vegetable Center, Taiwan, UK aid from the UK government, United States Agency for International Development (USAID), Australian Centre for International Agricultural Research (ACIAR), Germany, Thailand, Philippines, Korea, and Japan.

